# Top-down Mass Spectrometric Analysis of a Lipooligosaccharide from the Human Commensal *Bacteroides fragilis*

**DOI:** 10.1101/2024.04.13.589361

**Authors:** Tiandi Yang, Jason Daugherty, Dennis L. Kasper

## Abstract

Lipopolysaccharides (LPS) and lipooligosaccharides (LOS) are ubiquitous structures found on the outer membrane of gram-negative bacteria. *Bacteroides fragilis*, is a gram-negative anaerobe commonly inhabiting the human colon. The LOS of this organism is known to trigger a type I interferon response in dendritic cells. However, detailed structural analysis of this LOS has been largely elusive. Using top-down mass spectrometry, we have unraveled the comprehensive fine structure of *B. fragilis* LOS. Our analysis reveals that this LOS has a poly-galactose-rhamnose-KDO-lipid A architecture, which can be modified by hexuronic acid and ethanolamine via phosphodiester linkages. The lipid moiety typically includes three to five acyl chains of varying length on a glucosamine disaccharide. This investigation lays the groundwork for deeper immunological exploration of *B. fragilis* LOS and underscores the efficacy of top-down mass spectrometry in characterizing intact LOS/LPS structures and their modifications.

## Introduction

Lipopolysaccharides (LPS) are a group of glycolipids making up the outer leaflet of the asymmetrical outer membrane (OM) of Gram-negative bacteria. LPS generally contains three structural domains: the lipid A, which anchors the whole LPS to the OM; the O-antigen, which is the polysaccharide antigen located at the outer-most layer of gram-negative bacteria; and the core-oligosaccharide linking lipid A and the O-antigen.

Lipid A is essential for bacterial viability, and its biosynthesis and structure tend to be conserved through various organisms (*1*, *2*). The classical lipid A is a diphosphorylated glucosamine disaccharide modified by six acyl chains, and this glycolipid is a target of proinflammatory innate immune responses. Lipid A is recognized by the Toll-like receptor 4/myeloid differentiation factor 2 (TLR4/MD2) complex found on host cells (*3–6*), and the stimulation of TLR4/MD2 triggers the expression of cytokines crucial for the killing and clearance of pathogens. Cationic antimicrobial (*7*, *8*) peptides (CAMPs) can neutralize the phosphate groups, resulting in penetration of the LPS barrier surrounding the cell, ultimately leading to bacterial death. Some bacteria have evolved biosynthetic pathways resulting in extensive modifications of classical lipid A (*2*) allowing evasion of an inflammatory host response by becoming more tolerogenic. Such modifications include the reduction of acyl chain numbers (*9*), elongation of the acyl chain lengths (*10*, *11*), developing branched acyl chains (*12*, *13*), removal of one phosphate group (*14*, *15*), and blockage of the phosphates with positively charged groups (*10*, *16–21*). These modifications can significantly regulate proinflammatory immune responses (*6*), leading to a more tolerogenic life-long colonization as exemplified by organisms inhabiting the human gut.

The core-oligosaccharide of LPS almost always contains 3-deoxy-d-manno-octulosonic acid (KDO) as its reducing end sugar linking to lipid A, since KDO is added to lipid A during the conserved lipid A biosynthesis. A significant portion of characterized bacterial core oligosaccharides contain two KDOs and a few heptoses, but there are exceptions such as *Helicobacter pylori* (*21*, *22*), and *Bacteroides vulgatus* (*23*). Removal of one KDO was reported to help evade TLR2-mediated immune responses (*24*). The O-antigen is normally not essential for bacterial survival in culture, and it shows great structural diversity, typically serving as a target for antibodies directed to the microbe and evasion of the humoral immune system. When LPS does not carry the O-antigen, it is normally called rough-mode LPS or lipooligosaccharide (LOS).

*Bacteroides fragilis* (*B. fragilis*) is a gram-negative anaerobic bacterium commonly found in the human colon. It produces various surface capsular polysaccharides (CPS), LPS and LOS. Understanding the CPS structures has been a focus of previous research due to their immune activities, such as the ability to trigger T-cell-dependent immune responses (*25*). Although it is known that *Bacteroides* LOS induces a strong type I interferon response (*26*), the complete structure of the *Bacteroides* LOS is unknown. In this study, we present a nearly complete structure of *B. fragilis* LOS, determined through top-down and tandem mass spectrometry. Our findings reveal that the LOS has a poly-galactose-rhamnose-KDO-lipid A structure, which can be modified by hexuronic acid (HexA) and ethanolamine (EtN). Additionally, lipid A can carry three to five acyl chains with varying lengths. The polyhexose-deoxyhexose architecture and the lipid A structure are consistent with previous research (*27*, *28*). Our data highlight a heavily modified KDO-lipid A structure that was not previous reported. Our structural investigation paves the way for further immunological investigation of the glycolipid and establishes top-down mass spectrometry as a powerful tool to determine intact and modified LOS structures *de novo*.

## Material and Methods

All chemicals were purchased from Sigma-Aldrich unless otherwise indicated.

### LOS/PSA extraction and purification

*B. fragilis* bacteria were cultured using a minimal medium and a hot phenol/water extraction was performed as previously described with minor modifications (*29*). Briefly, the crude extract was washed with diethyl ether and treated by DNAse (Worthington Biochemical Corporation, US), RNAse (Worthington Biochemical Corporation, US), and Pronase K (Calbiochem, US). Polysaccharides were precipitated using 80% ethanol and the precipitate was solubilized in 1.5% deoxycholate buffer (pH=8), and the LOS/PSA was separated by a S400 column (2 m) equilibrated in 1.5% deoxycholate buffer. Deoxycholate was mostly removed by using tangential flow dialysis, and the resulting LOS solution was lyophilized.

### Mild acid hydrolysis

LOS was dissolved in a 100 mM sodium acetate buffer (pH=4.4) and incubated at 80 °C overnight. The lipid A and core-oligosaccharide were separated by a CHCl_3_/H_2_O extraction. The aqueous fraction containing the core-oligosaccharide was dialyzed and lyophilized. This protocol has been used before (*30–32*) and the author’s previous experiments indicate it is more efficient than acetic acid mild acid hydrolysis.

### Direct Infusion

A dilute (0.1-0.2 mg/mL) 0.1% formic acid (Fisher Scientific, US) solution of LPS was manually injected into the ESI source. All MS experiments were performed on a Q-Exactive Orbitrap mass analyzer (Thermo, US). The spray voltage was 3500 KV. The shealth gas flow rate was 50 µL/min and aux gas flow rate 15 µL/min. The capillary temperature was 300 °C. The S-lens RF level was 60%. MS was operated under the negative ion mode and the resolution was set to 70 K. AGC target was 1000000 and maximum injection time was 100 ms.

### HPLC-MS

The HPLC separation was performed using a Vanquish LC system (Thermo, US) equipped with a Kinetex C8 column (100 Å, 2.6 µm, 2.1*75 mm, Phenomenex, US) heated to 40 °C. Mobile phase A is pure water, B is isopropanol, and C is 2% formic acid. The mobile phase flow rate is 200 μL/min. Mobile phase C is always kept at 5% to provide formic acid. From 0 to 3.5 min, the B goes from 0% to 65%. From 3.5 to 10.5 min, B goes to 60% to 85%. The B was kept at 85% for 5 min before going down to 0% at 17 min. The B was kept at 0% for another 5 min before the end of the method. The MS was done on the same Orbitrap mass analyzer. The spray voltage was 3500 KV. The sheath gas flow rate was 50 µL/min and aux gas flow rate 15 µL/min. The capillary temperature was 300 °C. The S-lens RF level was 60%. MS was operated under the negative ion mode. For MS, the resolution was set to 70 K and mass range was 1000-3000 Da. Two micro-scans were accumulated to generate a single MS spectrum. AGC target was 3000000 and maximum injection time was 100 ms. For MSMS analysis, the resolution was 17.5 K. AGC target was 100000 and maximum injection time was 50 ms. Isolation windows was 4 Da.

### In-source CID

was performed for both direct infusion (for PSA) and HPLC-MS (for LOS) experiments. The fragmentation energy was set to 75 and 60 eV for PSA and LOS, respectively.

### NMR

The sample was subjected to H-D exchange before transferred to a 5 mM NMR tube. ^1^H, HSQC, and TOCSY spectra were recorded 298 K by a 400MHz Agilent NMR spectrometer.

### GC-MS monosaccharide and linkage analysis

Sample preparation for GC-MS monosaccharide (*33–35*) and linkage analysis (*36, 37*) are highly sophisticated, and we followed the protocol of a recent publication (*21*). The GC-MS experiments were done with either a Thermo GC-qExactive instrument equipped with a DB-5ms GC column (30 m, 0.25 mm, 0.25 µm) or an Agilent GC-MS equipped with a DB-624 column (30 m, 0.25 mm, 1.40 μm).

## Results and Discussion

### Direct infusion-based top-down MS

Mass spectrometry (MS) was usually utilized for characterizing intact LOS with well-annotated structures before (*38*, *39*). We extracted our sample from *B. fragilis delta 44*, which primarily produces a high-molecular-weight polysaccharide (PSA) and the LOS of interest in this study. Since the fine structure of the LOS is unknown, we started our research by manually infusing a dilute LOS solution into an electrospray (ES) ion source connected to an Orbitrap mass analyzer to test ionization efficiency. The LOS molecules were readily ionizable under a negative ion mode, and our analysis generated highly intricate spectra. (**Fig. 1a**). The most intense peaks are two pairs of MS signals at *m/z* 1264.53^2-^ and 1286.05^2-^ as well as 1391.65^2-^ and 1413.17^2-^ Da (depicted as red MS peaks in **Fig. 1**). We attributed the consistent 43.0 Da mass difference within the two pairs of MS peaks to an ethanolamine (EtN) modification on the LOS. Additionally, a 254.24 Da mass difference between the MS peaks at *m/z* 1264.53^2-^ and 1391.65^2-^, and 1286.05^2-^ and 1413.17^2-^ Da indicated that the addition of hydroxyhexadecanoic acid to the lower *m/z* ions generates the higher *m/z* ions. A further 254.24 Da addition to the peaks at *m/z* 1391.65^2-^ and 1413.17^2-^ Da ions produces peaks at *m/z* 1518.76^2-^ and 1540.29^2-^ (depicted as red MS peaks in **Fig. 1 a** and **d**). This implies that the lipid A part of the LOS can bear different numbers of acyl chains. Multiple satellite peaks are located around these highly intensive peaks with 14.0 Da mass differences (**Fig. 1 b** and **c**), indicating varying acyl chain lengths. Subsequent analysis (**Fig. 3** and **4**) strongly suggested that the lipid A portion can carry three to five acyl chains with differing carbon chain lengths. For instance, the MS peaks at *m/z* 1286.05^2-^, 1413.17^2-^, and 1540.29^2-^ Da corresponded to tri-, tetra-, and penta-acylated LOS, respectively. The tri- and tetra-acylated LOS MS signals dominated the spectrum, with only a low quantity of penta-acylated LOS observed. It remains uncertain whether the *delta 44* mutation influences the acylation status of lipid A, but lipid A tri-acylation is relatively uncommon.

**Figure 1.**
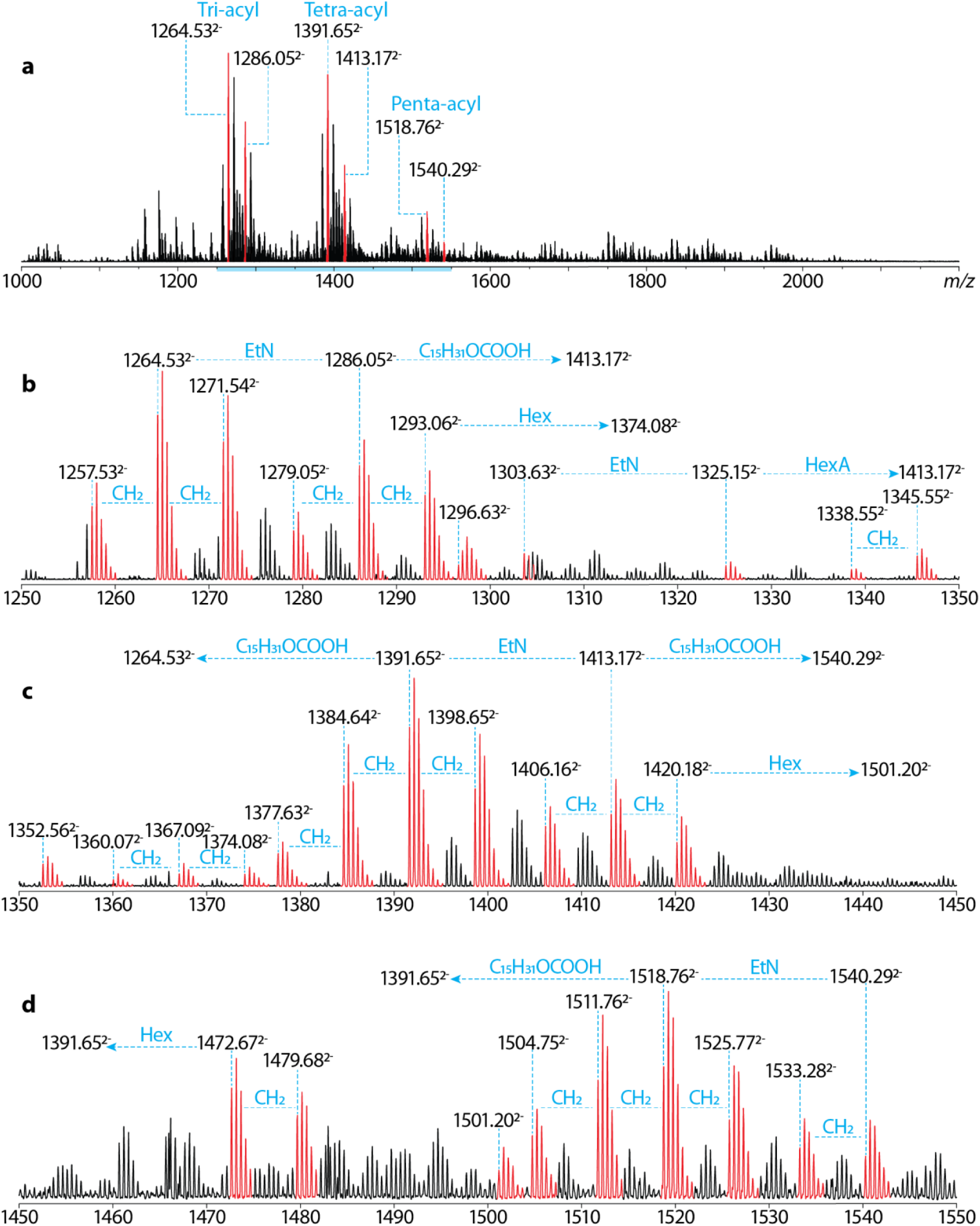
Direct infusion of intact LOS extracted from *B. Fragilis delta 44*. A dilute LOS solution was manually infused into the electrospray ion source, and a highly complicated MS profile (**a**) was collected by an orbitrap mass analyzer. The 1250-1550 Da mass range (the blue-shaded region) is zoomed in and shown in (**b**, 1250-1350 Da), (**c**, 1350 to 1450 Da), and (**d**, 1450-1550 Da). LOS is a glycolipid constructed by covalently linking oligosaccharide/short polysaccharide to lipid A. The spectrum indicates that the core-oligosaccharide is semi-conserved, whereas the lipid A carries tir-, tetra-, and penta-acyl chains with different lengths (see MS signals with CH_2_ mass difference). MS peaks with *m/z* differences corresponding to hexose (Hex), ethanolamine (EtN), and hexuronic acid (HexA) were observed. Further analysis (see Fig. 3 and 4) indicates Hex directly extending the core-oligosaccharide, while EtN and HexA modify LOS presumably via phosphodiester linkages. Tri-acylated LOS shows slightly higher intensity than tetra-acylated LOS and is significantly stronger than penta-acylated LOS.

### Top-down HPLC-MS

Since the direct infusion of intact LOS generated highly complex mass spectra, we separated the LOS by reverse-phase high-performance liquid chromatography (RP-HPLC) before in-line MS analysis to reduce the complexity of the MS spectra and decrease the sample usage to a low microgram level. As expected, the LOS was separated based on hydrophobicity: the tri-acylated LOS peak was eluted first and slightly more intense than the following tetra-acylated LOS peak (**Fig. 2 a-c**). The penta-acylated LOS eluted at the latest with a broad LC peak (**data not shown**), indicating an over-optimal interaction between the column and the LOS, which is resolvable by using chloroform to elute the column (*39*). However, we adhered to the current isopropanol protocol to mitigate toxicity concerns in an open laboratory setting. This protocol proved sufficient for separating tri- and tetra-acylated LOS for MS analysis (**Fig. 2 b** and **c**). We accumulated all MS spectra collected during the 10.94- and 12.16-min LC peaks, which showed the same poly-hexose addition pattern within both LC peaks, with the fourth MS signals in both clusters of hexose (Hex) addition possessing the dominant intensities (**Fig. 2 d** and **e**). This indicated that LOS with different acyl chains shares a very similar, if not identical, polyHex-containing core-oligosaccharide architecture. Therefore, we focused our further analysis on the tri-acylated LOS, which is characterizable by the *m/z* 1286.05^2-^ Da peak-containing MS peak cluster. This is also due to their lower molecular weight and higher MS intensities. The tetra-acylated LOS has an additional disadvantage as some of its MS signals interfere with the following-up in-source fragmentation-based MS3 analysis. For instance, tetra-acylated LOS and its lipid A fragment can both have *m/z* around 1420.

**Figure 2.**
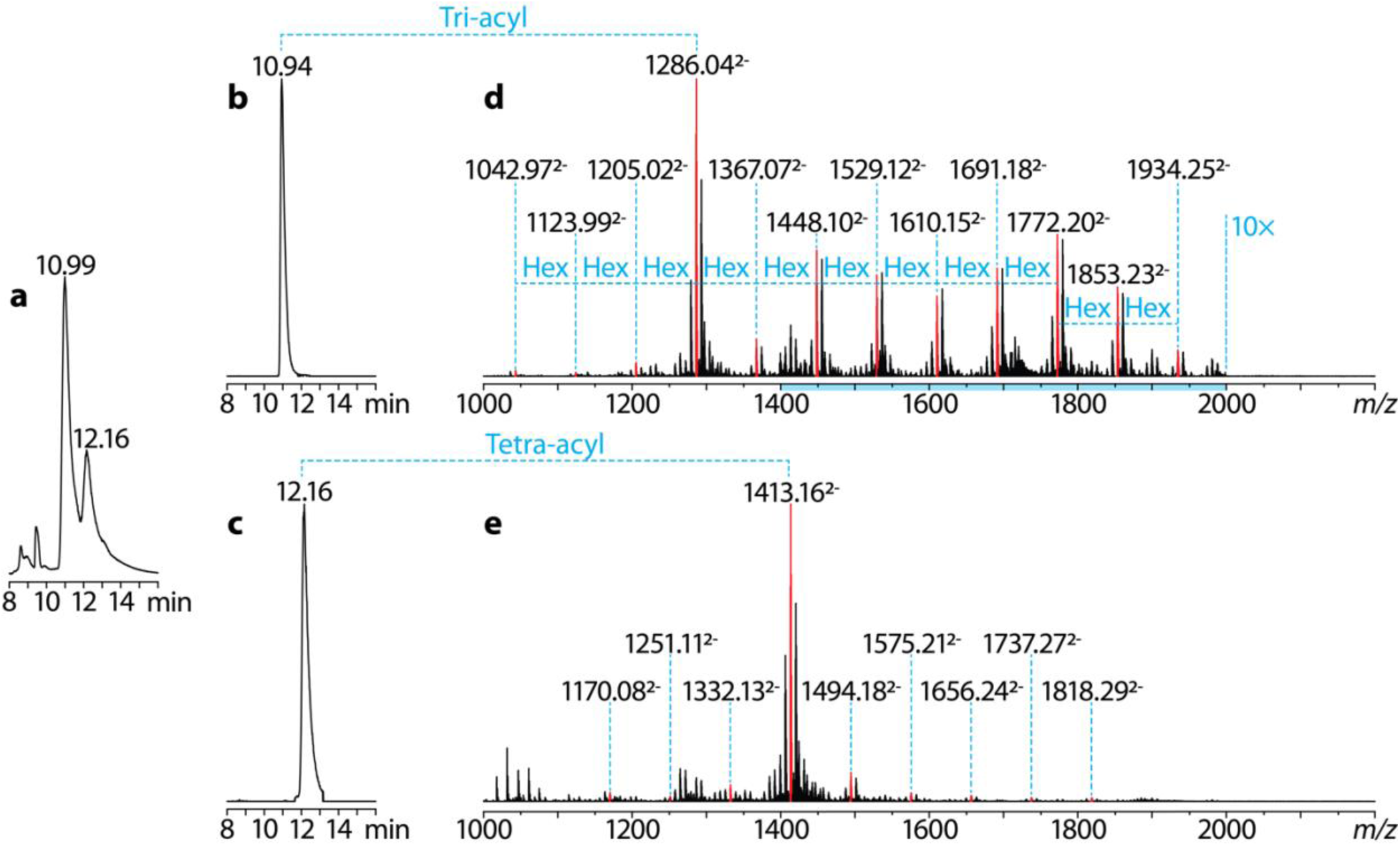
HPLC-ESI-MS analysis of intact LOS extracted from *B. fragilis delta 44*. Intact LOS was separated by reverse-phase HPLC columns before inline analyzed by an ESI-orbitrap mass spectrometer. The total ion chromatogram is shown in (**a**). The extracted ion chromatograms of selected tri-acylated and tetra-acylated LOS are shown in (**b**) and (**c**), respectively. Accumulated MS spectra of 10.94- and 12.16-min LC peaks are shown in (**d**) and (**e**), respectively. Selected MS signals were colored red and annotated with *m/z* values, as these strong MS peaks are in the clusters of peaks with a *m/z* difference corresponding to Hex. Most unannotated MS peaks correspond to the “same structure” with different acyl chain lengths, and the same clusters can also be observed for the unannotated peaks. Intensities of the MS peaks within the mass range of 1400-2000 Da (blue-shaded region) in (**d**) were amplified ten times, whereas (**e**) was left untouched to facilitate the comparison of MS peak intensities. MS peaks at *m/z* 1286.04 and 1413.16 Da dominate their correspondent spectra. Further analysis reveals both MS peaks corresponding to LOS containing three Hex residues (see Fig. 3).

### MSMS

All MS signals in the *m/z* 1286.05^2-^ Da peak cluster (**Fig. 2d**) were further analyzed by MSMS (**Fig. 3**). The higher-energy collisional dissociation (HCD) effectively fragmented the LOS molecular ions, and as anticipated, their MSMS spectra shared certain ions corresponding to the same lipid A and LOS modification (green dot lines in **Fig. 3**, **Fig. 4**). Specifically, the fragment ion at *m/z* 1165.8 Da corresponds to tri-acylated mono-phosphorylated lipid A, which can lose a pentadecanoic acid to form the ion at *m/z* 923.6 Da. In addition, there is a 176.0 Da structure adding to the lipid A, giving rise to another ion at *m/z* 1341.71 Da. We also noticed a strong MSMS peak at *m/z* 273.00 Da, potentially corresponding to a phosphorylated hexuronic acid (HexA-P). These observations indicated lipid A is modified by a hexuronic acid (176.04 Da) via a phosphodiester bond. More importantly, this lipid A seems to be conserved, while all the differences between the LOS structures were attributed to the different lengths of polyHex linking to the lipid A (blue arrows in **Fig. 3a**). The polyHex chain length ranges from 1-11 monosaccharide units, and it directly links to an ion at *m/z* 426.12. We subjected this ion to in-source fragmentation-based MS3 analysis (see below) and this ion corresponds to a deoxy-hexose addition (**Fig. 3b**) to an *m/z* 262.05 Da ion that is most likely associated with 3-Deoxy-d-manno-octulosonic acid (KDO). Gas chromatography (GC)-MS monosaccharide analysis (**sFig.1 a** and **b**) indicates the hexose is mainly galactose (Gal), and the deoxy-hexose is rhamnose (Rha). Linkage analysis indicates the hexose is 6- and 4,6-linked and the deoxy-hexose is 2-linked (**sFig.1 c**-**e**).

**Figure 3.**
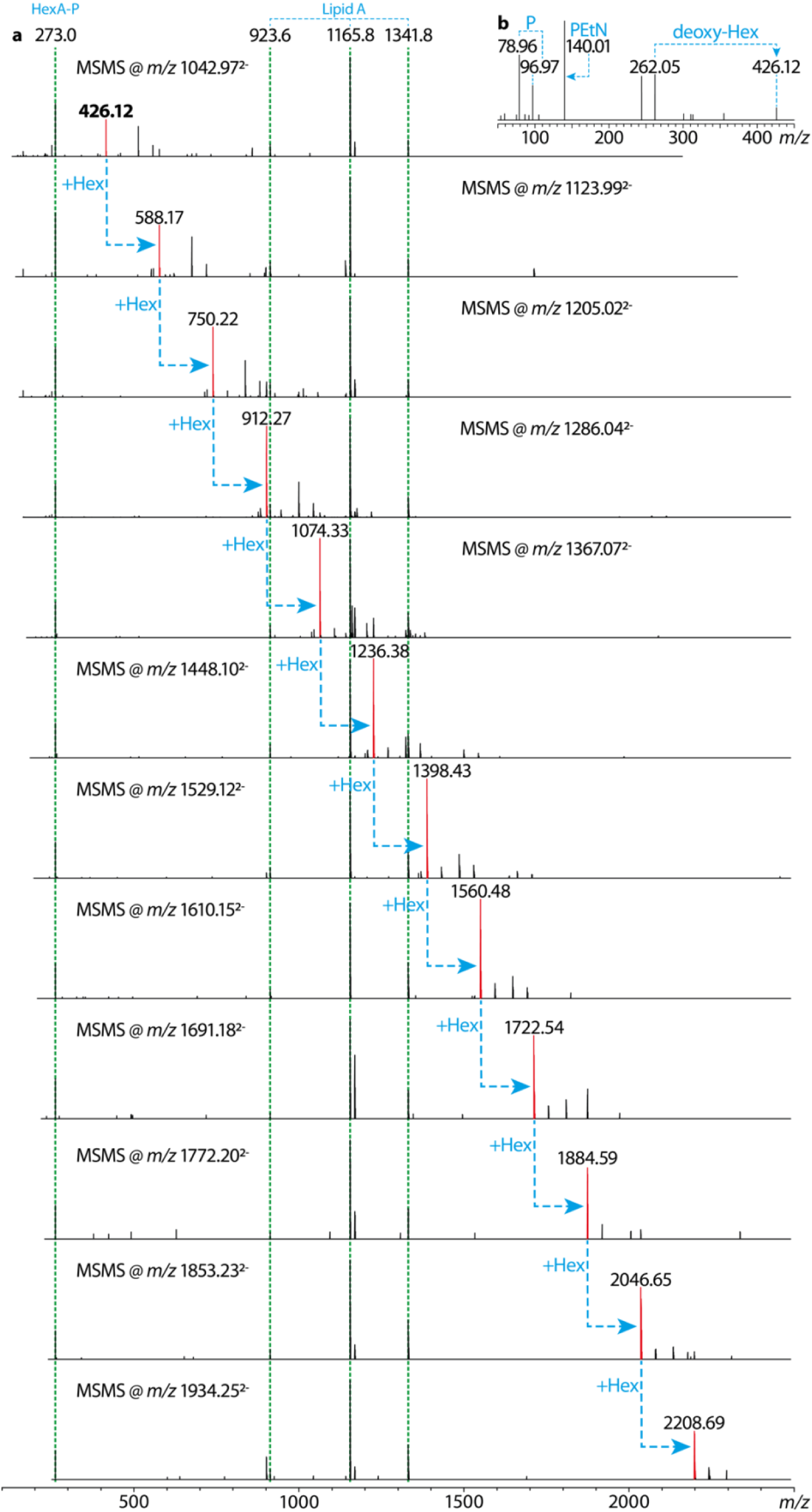
HCD MSMS analysis of tri-acylated LOS. All selected peaks in Fig. 2 were further analyzed by HCD-based MSMS analysis (**a**). All the MSMS spectra share the same MS peaks at *m/z* 273.0, 923.6, 1165.8 and 1341.8 Da. There are very minor differences (0.01 Da level) in the mass accuracy of the MSMS spectra, so *m/z* values with only one decimal number were reported. The peak at *m/z* 273.0 Da corresponds to HexA-phosphate (HexA-P), whereas the three remaining peaks are from lipid A and its fragments (see Fig. 4 and 5). We found a Hex addition pattern to an MSMS peak at *m/z* 426.12 Da. This ion was subjected to in-source fragmentation-based MS3 analysis, and the spectrum strongly indicated the ion contains deoxy-hexose (deoxy-Hex) and phosphoethanolamine (PEtN). These MSMS and MS3 data revealed a PEtN-modified poly-hexose-deoxyhexose core-oligosaccharide architecture.

Overall, the data indicated the dominant peak at *m/z* 1286.05^2-^ Da corresponds to a branched Gal_3_- Rha-KDO oligosaccharide linking to a tri-acylated monophosphorylated lipid A, which is further modifiable by HexA and PEtN. However, the direct addition of these residues does not match the observed *m/z* value. More importantly, the *m/z* 262.05 Da ion does not directly match the mass of KDO fragment ions. So, we next focused on MSMS of the *m/z* 1264.53^2-^ and 1286.05^2-^ Da ions (see **Fig.1 a** and **b**, **Fig. 4a**), and a more careful interrogation of the MSMS data revealed a fragment ion potentially corresponding to intact core-oligosaccharide at *m/z* 1230.27 Da (**Fig. 4b**). This ion seems to be related to a few ions at *m/z* 912.27, 956.26, and 1054.24 Da. For instance, the mass difference between ions at *m/z* 1230.27 and 1054.24 Da is 176.03 Da corresponding to HexA. The *m/z* 1054.24 Da can lose phosphate, forming the *m/z* 965.27 Da ion that can further decarboxylate (minus 43.99 Da) into the abundant ion at *m/z* 912.27 Da. This indicates that the core-oligosaccharide could carry at least another phosphate and a HexA group.

**Figure 4.**
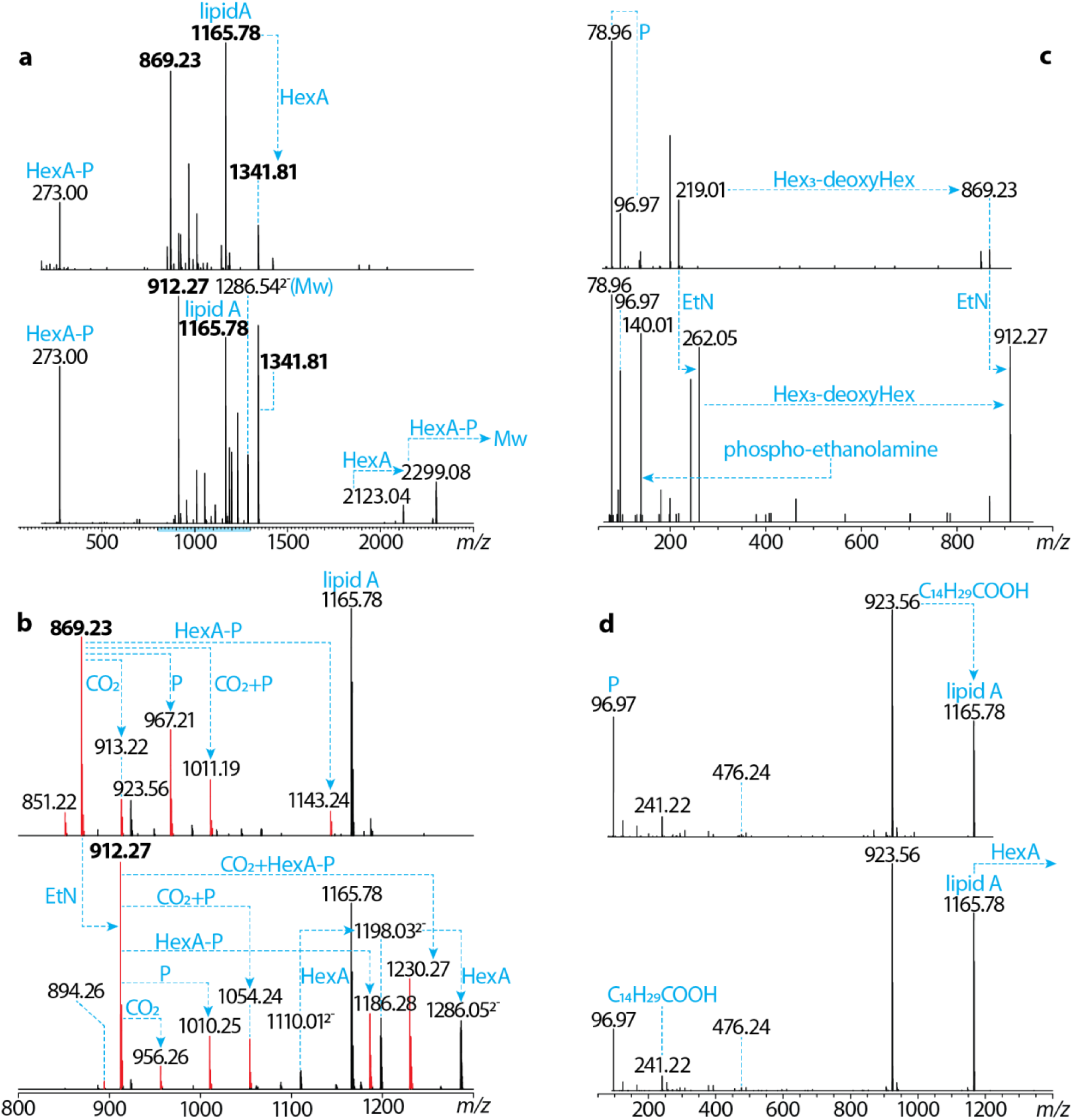
HCD analysis of tri-acylated LOS containing three Hex residues. (**a**) HCD MSMS spectra of MS peak at *m/z* 1264.53 (upper panel) and 1286.05 (lower panel) Da (see Fig. 1b). HCD generated the fragment ions corresponding to HexA-P (*m/z* 273.00 Da), lipid A (*m/z* 1165.78 and 1341.81 Da), and core-oligosaccharide (*m/z* 869.23 and 912.27 Da). The MSMS spectra were zoomed into the 800-1300 Da mass range and shown in (**b**). The same addition pattern (the addition of CO_2_, P, CO_2_+P, and HexA-P) was observed for both *m/z* 869.23 and 912.27 Da peaks, indicating a HexA-P modification on the core-oligosaccharide. The lipid A and core-oligosaccharide fragments were further analyzed by in-source fragmentation-based MS3 analysis. (**c**) MS3 analysis of *m/z* 869.23 (upper panel) and 912.27 (lower panel) Da. In the lower panel, a PEtN ion was observed at *m/z* 140.01 Da. The elimination of deoxyHex-Hex3 from 912.29 resulted in generating an ion at *m/z* 262.05 Da. The same ion was also observed in the MS3 spectrum of *m/z* 426.12 Da (see Fig. 3b). In the upper panel, the PEtN ion was missing and the *m/z* 262.05 Da peak shifted to *m/z* 219.01 Da, with a difference corresponding to EtN. These data further indicate the core-oligosaccharide has a poly-hexose-deoxyhexose architecture modifiable by PEtN. (**d**) MS3 analysis of *m/z* 1165.78 (upper panel) and 1341.81 (lower panel) Da peaks. The two spectra are nearly identical, indicating there is a HexA addition to the tri-acylated lipid A.

### In-source fragmentation-based MS3

To further interrogate the structure of the core-oligosaccharide, we performed in-source fragmentation-based MS3 on the core-oligosaccharide, especially for the ions at *m/z* 869.23 and 912.27 Da (**Fig. 4c**). Briefly, if we directly fragment molecular ions at the ion source (in-source fragmentation), the scanned MS spectra effectively record MSMS information, while the MSMS spectra record MS3 information. The two mass spectra share the same phosphate fragment ion and the pattern of eliminating a Hex3-deoxyHex tetra-saccharide. Their mass difference is due to the EtN modification found on the latter ions. The *m/z* 912.27 Da ion was fragmented into an ion with *m/z* of 140.01 Da, which perfectly matches the mass of phosphoethanolamine. This confirms the EtN links on a phosphate on the core-oligosaccharide. In addition, the tetra-saccharide elimination of *m/z* 869.23 and 912.27 Da ion leads to an ion at *m/z* 219.01 Da and the previously observed 262.05 Da ion, respectively. Their mass difference matches the mass of EtN, which reveals the PEtN modification locates on the KDO-associated ion. We next analyzed the aqueous fraction of the mild acid hydrolyzed LOS (**sFig.1 f** and **g**) and found an ion at *m/z* 974.28 Da corresponding to the same structure as the *m/z* 956.27 Da ion. The 18.0 Da mass shift is due to the difference between fragmentation and hydrolysis. The fragment ion at *m/z* 140.01 Da was detected in the MSMS spectra of the *m/z* 974.28 Da ion as well, further supporting the core-oligosaccharide modifiable by PEtN. We then performed NMR spectroscopy and found characteristic ^1^H and ^13^C chemical shifts (*40*) of PEtN in the HSQC and TOCSY spectra (**sFig.1 h** and **i**). Unfortunately, we were not able to observe the core-oligosaccharide modified by HexA after the hydrolysis. Collectively, the MSMS and in-source fragmentation-based MS3 data reveal the KDO residue in the core-oligosaccharide is heavily modified by both EtN and HexA via two phosphodiester bonds.

The in-source fragmentation-based MS3 also gave us more insight into the structure of lipid A. A strong MSMS peak at *m/z* 241.22 Da corresponding to pentadecanoic acid was found. This acyl chain could link to a hydroxyl group. A low MSMS peak at *m/z* 476.24 Da corresponds to a lipid A fragment, indicating the phosphorylated GlcN linking to a C16 N-acyl chain. This ion also implies the mono-phosphate links on the reducing end of the GlcN (1-position). The remaining C17 N-acyl chain might link to another GlcN. The fragmentation pattern of *m/z* 1165.79 and 1341.82 are identical, and their mass difference corresponds to a HexA, which further confirms the modification on lipid A.

After successfully implementing the technique for analyzing intact LOS molecules, we performed a manual infusion of *B. fragilis* PSA (*41*) upon recognizing the potential of the in-source fragmentation for studying glycoconjugates carrying long polysaccharides. It was impossible to annotate the collected MS spectra (**data not shown**) due to their complexity and low quality without in-source fragmentation. However, upon employing in-source fragmentation, we were able to readily “sequence” the glycan repeats (**Fig. 6**). Consequently, the application of our approach seems to extend beyond LOS characterization and is readily transferrable to longer polysaccharides and LPSs.

**Figure 5.**
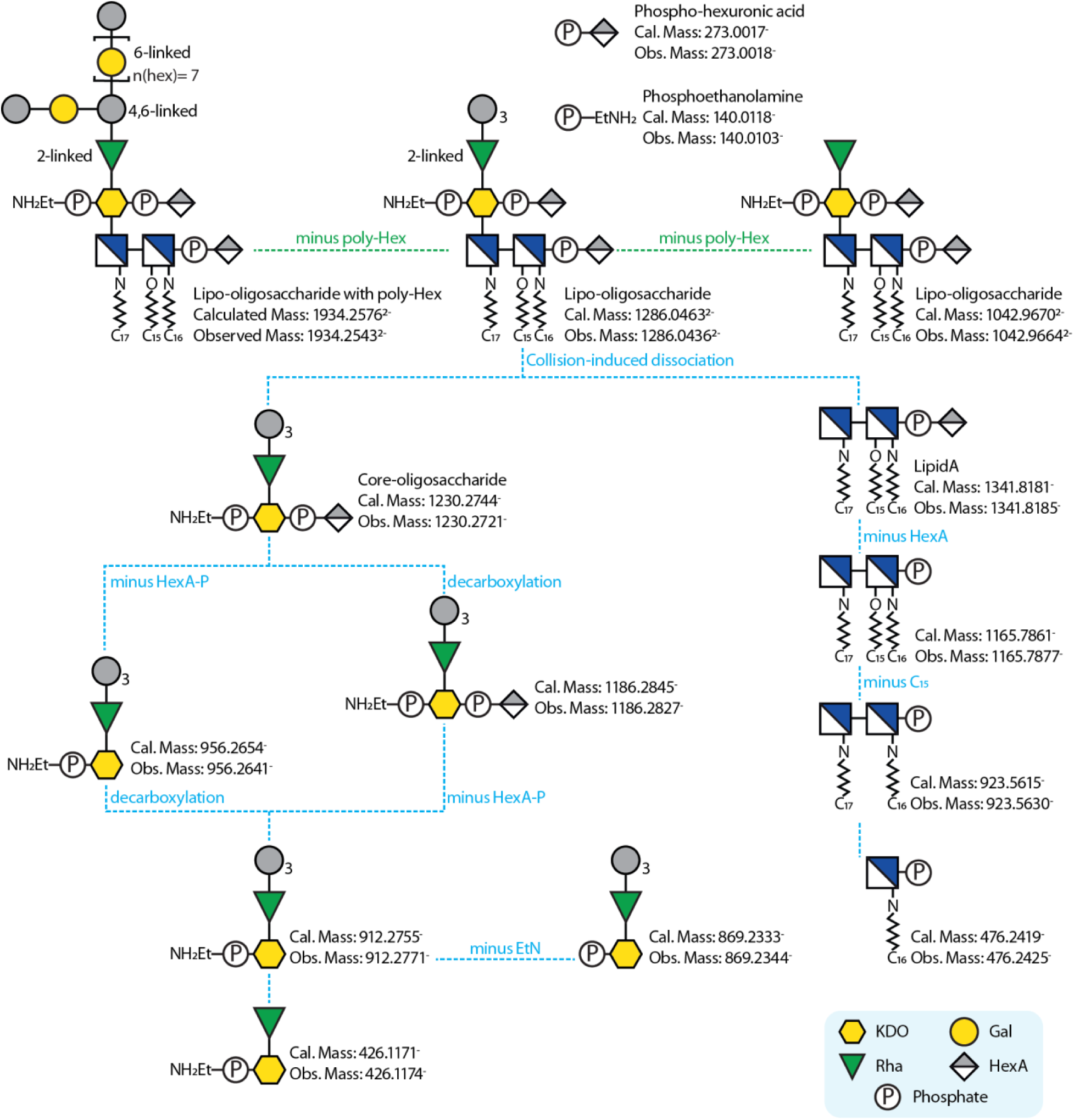
Fragmentation pathway of tri-acylated LOS. This specific LOS is constructed by linking a poly-Hex-deoxy-Hex-KDO oligo-/poly-saccharide to a tri-acylated lipid A (green dot line connected). The tri-acylated LOS containing three Hex residues was subjected to HCD fragmentations (blue dot line connected). Important fragments were listed with calculated and observed masses. We are working to determine the exact linkages and the stereochemical properties of the monosaccharides, but existing data indicate most Hex residues are galactose, and the deoxy-Hex is a 2-linked rhamnose. There is a 4,6 linked-Hex, indicating the glycan branches at least once. The monophosphate on the lipid A links at the reducing end of the glucosamine, and it likely links to a HexA. PEtN most likely links to the KDO directly. HexA-P modifies both the core-oligosaccharide and lipid A. It might connect to KDO via a phosphodiester linkage, but we cannot exclude the possibility of glycosidic or other linkages.

**Figure 6.**
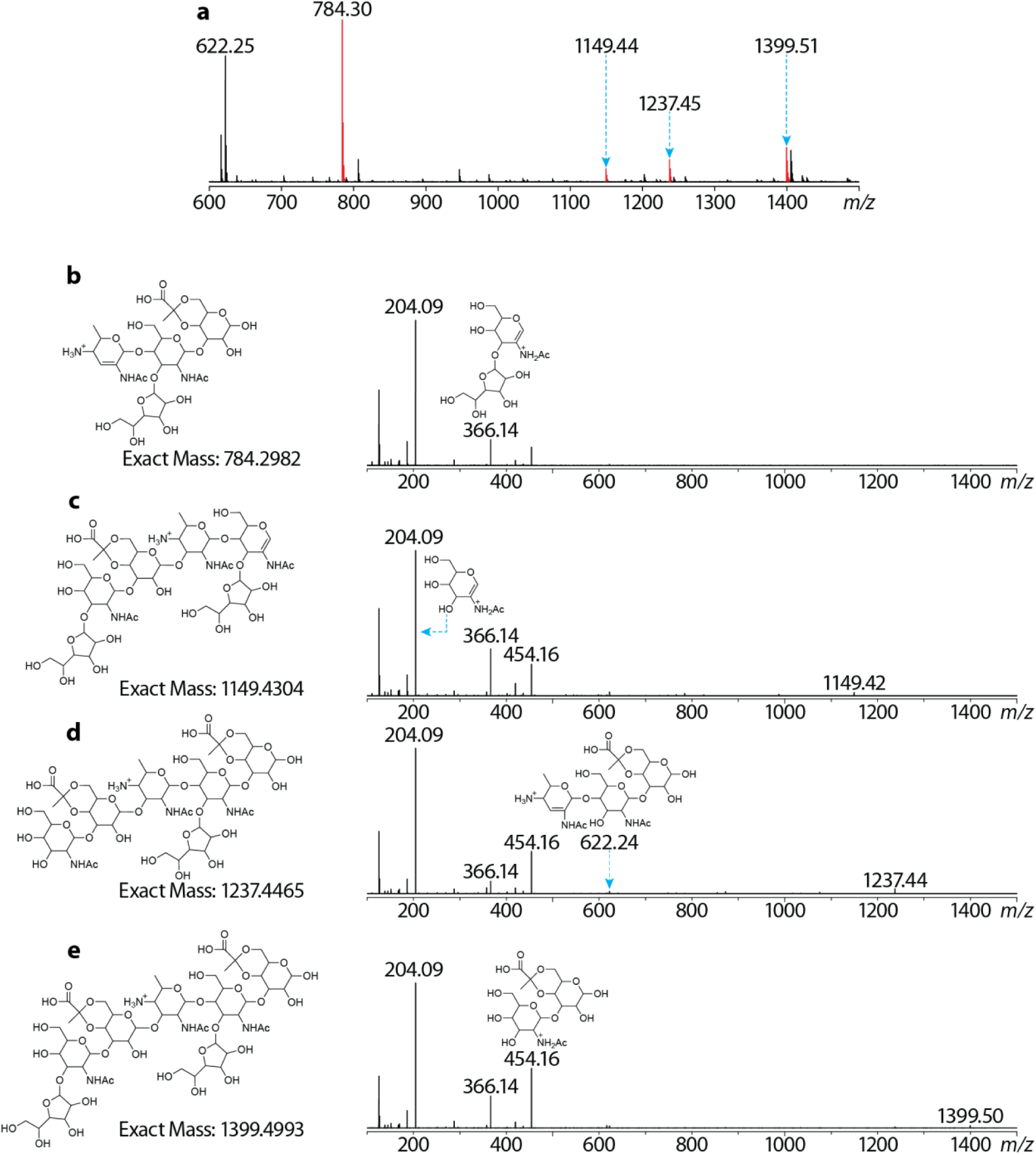
MS In-source fragmentation characterization of intact *B. fragilis* PSA. (**a**) MS scan of in-source fragmented PSA. A few MS signals corresponding to the PSA fragments are annotated with *m/z*. The red peaks were further characterized by MSMS (in-source fragmentation-based MS3). (**b-e**) In-source fragmented MS3 of PSA. The proposed structures are annotated with *m/z* on the left of the corresponding MSMS spectra. Note that these MSMS data do not provide stereochemical information, but the fragmentation indicates the pyruvate-galactose as the non-reducing end monosaccharide (**b**).

In summary, our structural analysis indicates LOS extracted from the *B. fragilis delta 44* strain is a complex mixture of glycolipids. Both the glycan and lipid A parts contribute to structural diversity. The core oligosaccharide is a polyHex-deoxyHex-KDO glycan mixture with potential HexA-P and PEtN modifications, while the lipid A carries three to five acyl chains with variable lengths and modifiable by HexA. This work addresses the central role of mass spectrometry glycoconjugate analysis: even an entry-level Orbitrap mass spectrometer can carry out highly sophisticated glycolipid analysis.

The so-called core-oligosaccharide contains a galactose polymer, and we cannot exclude the possibility that this polymer is the O-antigen part of a short LPS. This question needs to be resolved by further biosynthetic investigations. Meanwhile, we are purifying samples further to facilitate NMR analysis so that we can confirm the conclusions based on our MS data and assign all the remaining stereochemical features.

## Acknowledgements

We would like to acknowledge Byoungsook Goh and Sungwhan F. Oh for the maintenance of the LC-MS instrument and their useful suggestions. We thank Harvard Center for Mass Spectrometry and Prof. Sloan Devlin’s lab for providing GC-MS platforms.

## Author contributions

TY and DLK conceived the idea. TY designed and carried out the experiments and data analysis. JD purified LOS and PSA samples. TY, DLK and JD wrote the manuscript.

## References

1. C. R. H. Raetz, C. Whitfield, LIPOPOLYSACCHARIDE ENDOTOXINS. Annu. Rev. Biochem. 71, 635–700 (2002).

2. C. R. H. Raetz, C. M. Reynolds, M. S. Trent, R. E. Bishop, Lipid A Modification Systems in Gram-Negative Bacteria. Annu. Rev. Biochem. 76, 295–329 (2007).

3. B. Lemaitre, E. Nicolas, L. Michaut, J. M. Reichhart, J. A. Hoffmann, The dorsoventral regulatory gene cassette spätzle/Toll/cactus controls the potent antifungal response in Drosophila adults. Cell 86, 973–983 (1996).

4. S. T. Qureshi, L. Larivière, G. Leveque, S. Clermont, K. J. Moore, P. Gros, D. Malo, Endotoxin-tolerant Mice Have Mutations in Toll-like Receptor 4 (Tlr4). J. Exp. Med. 189, 615– 625 (1999).

5. A. Poltorak, X. He, I. Smirnova, M.-Y. Liu, C. V. Huffel, X. Du, D. Birdwell, E. Alejos, M. Silva, C. Galanos, M. Freudenberg, P. Ricciardi-Castagnoli, B. Layton, B. Beutler, Defective LPS Signaling in C3H/HeJ and C57BL/10ScCr Mice: Mutations in *Tlr4* Gene. Science 282, 2085–2088 (1998).

6. B. S. Park, D. H. Song, H. M. Kim, B.-S. Choi, H. Lee, J.-O. Lee, The structural basis of lipopolysaccharide recognition by the TLR4-MD-2 complex. Nature 458, 1191–1195 (2009).

7. M. Vaara, T. Vaara, Polycations as outer membrane-disorganizing agents. Antimicrob. Agents Chemother. 24, 114–122 (1983).

8. M. Vaara, T. Vaara, Outer Membrane Permeability Barrier Disruption by Polymyxin in Polymyxin-Susceptible and -Resistant Salmonella typhimurium. Antimicrob. Agents Chemother. 19, 578–583 (1981).

9. K. Kawahara, H. Tsukano, H. Watanabe, B. Lindner, M. Matsuura, Modification of the Structure and Activity of Lipid A in Yersinia pestis Lipopolysaccharide by Growth Temperature. Infect. Immun. 70, 4092–4098 (2002).

10. C. M. Stead, J. Zhao, C. R. H. Raetz, M. S. Trent, Removal of the outer Kdo from Helicobacter pylori lipopolysaccharide and its impact on the bacterial surface. Molecular Microbiology 78, 837–852 (2010).

11. C. M. Stead, A. Beasley, R. J. Cotter, M. S. Trent, Deciphering the Unusual Acylation Pattern of Helicobacter pylori Lipid A. J. Bacteriol. 190, 7012–7021 (2008).

12. R. A. Reife, S. R. Coats, M. Al-Qutub, D. M. Dixon, P. A. Braham, R. J. Billharz, W. N. Howald, R. P. Darveau, Porphyromonas gingivalis lipopolysaccharide lipid A heterogeneity: differential activities of tetra- and penta-acylated lipid A structures on E-selectin expression and TLR4 recognition. Cell. Microbiol. 8, 857–868 (2006).

13. S. R. Coats, A. B. Berezow, T. T. To, S. Jain, B. W. Bainbridge, K. P. Banani, R. P. Darveau, The Lipid A Phosphate Position Determines Differential Host Toll-Like Receptor 4 Responses to Phylogenetically Related Symbiotic and Pathogenic Bacteria. Infect. Immun. 79, 203–210 (2011).

14. A. X. Tran, J. D. Whittimore, P. B. Wyrick, S. C. McGrath, R. J. Cotter, M. S. Trent, The Lipid A 1-Phosphatase of Helicobacter pylori Is Required for Resistance to the Antimicrobial Peptide Polymyxin. J. Bacteriol. 188, 4531–4541 (2006).

15. A. X. Tran, M. J. Karbarz, X. Wang, C. R. H. Raetz, S. C. McGrath, R. J. Cotter, M. S. Trent, Periplasmic cleavage and modification of the 1-phosphate group of Helicobacter pylori lipid A. Journal of Biological Chemistry 279, 55780–55791 (2004).

16. K. Nummila, Ii. Kilpeläinen, U. Zähringer, M. Vaara, Ii. M. Helander, Lipopolysaccharides of polymyxin B-resistant mutants of Escherichia coii are extensively substituted by 2-aminoethyl pyrophosphate and contain aminoarabinose in lipid A. Mol. Microbiol. 16, 271–278 (1995).

17. B. A. Scarbrough, C. R. Eade, A. J. Reid, T. C. Williams, J. M. Troutman, Lipopolysaccharide Is a 4-Aminoarabinose Donor to Exogenous Polyisoprenyl Phosphates through the Reverse Reaction of the Enzyme ArnT. ACS Omega 6, 25729–25741 (2021).

18. A. Anandan, G. L. Evans, K. Condic-Jurkic, M. L. O’Mara, C. M. John, N. J. Phillips, G. A. Jarvis, S. S. Wills, K. A. Stubbs, I. Moraes, C. M. Kahler, A. Vrielink, Structure of a lipid A phosphoethanolamine transferase suggests how conformational changes govern substrate binding. Proc. Natl. Acad. Sci. 114, 2218–2223 (2017).

19. E. Altman, V. Chandan, J. Li, E. Vinogradov, Lipopolysaccharide structure of Helicobacter pylori serogroup O:3. Carbohydrate Research 378, 139–143 (2013).

20. E. Altman, V. Chandan, J. Li, E. Vinogradov, Lipopolysaccharide structures of Helicobacter pylori wild-type strain 26695 and 26695 HP0826::Kan mutant devoid of the O-chain polysaccharide component. Carbohydrate Research 346, 2437–2444 (2011).

21. H. Li, T. Yang, T. Liao, A. W. Debowski, H.-O. Nilsson, A. Fulurija, S. M. Haslam, B. Mulloy, A. Dell, K. A. Stubbs, B. J. Marshall, M. Benghezal, The redefinition of Helicobacter pylori lipopolysaccharide O-antigen and core-oligosaccharide domains. PLoS Pathogens 13, e1006280 (2017).

22. A. P. Moran, B. Shiberu, J. A. Ferris, Y. A. Knirel, S. N. Senchenkova, A. V. Perepelov, P.- E. Jansson, J. B. Goldberg, Role of Helicobacter pylori rfaJ genes (HP0159 and HP1416) in lipopolysaccharide synthesis. FEMS microbiology letters 241, 57–65 (2004).

23. F. D. Lorenzo, M. D. Pither, M. Martufi, I. Scarinci, J. Guzmán-Caldentey, E. Łakomiec, W. Jachymek, S. C. M. Bruijns, S. M. Santamaría, J.-S. Frick, Y. van Kooyk, F. Chiodo, A. Silipo, M. L. Bernardini, A. Molinaro, Pairing Bacteroides vulgatus LPS Structure with Its Immunomodulatory Effects on Human Cellular Models. Acs Central Sci 6, 1602–1616 (2020).

24. N. A. Okan, S. Chalabaev, T.-H. Kim, A. Fink, R. A. Ross, D. L. Kasper, Kdo Hydrolase Is Required for Francisella tularensis Virulence and Evasion of TLR2-Mediated Innate Immunity. mBio 4, e00638–12 (2013).

25. E. B. Troy, D. L. Kasper, Beneficial effects of Bacteroides fragilis polysaccharides on the immune system. Front. Biosci. 15, 25 (2010).

26. K. L. Stefan, M. V. Kim, A. Iwasaki, D. L. Kasper, Commensal Microbiota Modulation of Natural Resistance to Virus Infection. Cell 183, 1312–1324.e10 (2020).

27. A. Weintraub, U. Zähringer, H. Wollenweber, U. Seydel, E. Th. Rietschel, Structural characterization of the lipid A component of Bacteroides fragilis strain NCTC 9343 lipopolysaccharide. Eur. J. Biochem. 183, 425–431 (1989).

28. A. Weintraub, U. Zähringer, A. A. Lindberg, Structural studies of the polysaccharide part of the cell wall lipopolysaccharide from Bacteroides fragilis NCTC 9343. Eur. J. Biochem. 151, 657–661 (1985).

29. A. Pantosti, A. O. Tzianabos, A. B. Onderdonk, D. L. Kasper, Immunochemical characterization of two surface polysaccharides of Bacteroides fragilis. Infect. Immun. 59, 2075– 2082 (1991).

30. J. C. Henderson, J. P. O’Brien, J. S. Brodbelt, M. S. Trent, Isolation and Chemical Characterization of Lipid A from Gram-negative Bacteria. J. Vis. Exp., e50623 (2013).

31. A. Tirsoaga, A. E. Hamidi, M. B. Perry, M. Caroff, A. Novikov, A rapid, small-scale procedure for the structural characterization of lipid A applied to Citrobacter and Bordetella strains: discovery of a new structural element. J. Lipid Res. 48, 2419–2427 (2007).

32. M. R. Rosner, J. Tang, I. Barzilay, H. G. Khorana, Structure of the lipopolysaccharide from an Escherichia coli heptose-less mutant. I. Chemical degradations and identification of products. J. Biol. Chem. 254, 5906–5917 (1979).

33. C. C. Sweeley, R. Bentley, M. Makita, W. W. Wells, Gas-liquid chromatography of trimethylsilyl derivatives of sugars and related substances. J. Am. Chem. Soc. 85, 2497–2507 (1963).

34. D. C. DeJongh, T. Radford, J. D. Hribar, S. Hanessian, M. Bieber, G. Dawson, C. C. Sweeley, Analysis of trimethylsilyl derivatives of carbohydrates by gas chromatography and mass spectrometry. Journal of American Chemical Society 91, 1728–1740 (1969).

35. J. F. Kennedy, S. M. Robertson, Mass-spectrometric fragmentation pathways of the O-trimethylsilyl derivatives of hexuronic acids and their lactones. Carbohydrate Research 67, 1–15 (1978).

36. S. Hakomori, A RAPID PERMETHYLATION OF GLYCOLIPID, AND POLYSACCHARIDE CATALYZED BY METHYLSULFINYL CARBANION IN DIMETHYL SULFOXIDE. Journal of Biochemistry 55, 205–208 (1964).

37. W. S. York, A. G. Darvill, M. McNeil, T. T. Stevenson, P. Albersheim, Isolation and characterization of plant cell walls and cell wall components. Methods Enzym. 118, 3–40 (1986).

38. B. L. Oyler, M. M. Khan, D. F. Smith, E. M. Harberts, D. P. A. Kilgour, R. K. Ernst, A. S. Cross, D. R. Goodlett, Top Down Tandem Mass Spectrometric Analysis of a Chemically Modified Rough-Type Lipopolysaccharide Vaccine Candidate. J. Am. Soc. Mass Spectrom. 29, 1221–1229 (2018).

39. D. R. Klein, M. J. Powers, M. S. Trent, J. S. Brodbelt, Top-Down Characterization of Lipooligosaccharides from Antibiotic-Resistant Bacteria. Anal. Chem. 91, 9608–9615 (2019).

40. U. Zähringer, S. Ittig, B. Lindner, H. Moll, U. Schombel, N. Gisch, G. R. Cornelis, NMR-based structural analysis of the complete rough-type lipopolysaccharide isolated from Capnocytophaga canimorsus. Journal of Biological Chemistry 289, 23963–23976 (2014).

41. A. O. Tzianabos, R. W. Finberg, Y. Wang, M. Chan, A. B. Onderdonk, H. J. Jennings, D. L. Kasper, T Cells Activated by Zwitterionic Molecules Prevent Abscesses Induced by Pathogenic Bacteria*. J. Biol. Chem. 275, 6733–6740 (2000).

